# HfaE is a component of the holdfast anchor complex that tethers the holdfast adhesin to the cell envelope

**DOI:** 10.1101/2022.07.20.500906

**Authors:** Nelson K. Chepkwony, Gail G. Hardy, Yves V. Brun

**Affiliations:** Département de microbiologie, infectiologie et immunologie, Université de Montréal, Montréal, Québec, Canada; Department of Biology, Indiana University, Bloomington, Indiana, USA

**Keywords:** Holdfast, Bacterial adhesin, Adhesion, Biofilm, Extracellular Polysaccharides, Caulobacterales.

## Abstract

Bacteria use adhesins to colonize different surfaces and form biofilms. The species of the Caulobacterales order use a polar adhesin called holdfast, composed of polysaccharides, proteins, and DNA to irreversibly adhere to surfaces. In *C. crescentus,* a freshwater Caulobacterales, the holdfast is anchored at the cell pole via the <u>h</u>oldfast <u>a</u>nchor (Hfa) proteins HfaA, HfaB, and HfaD. HfaA and HfaD co-localize with holdfast and are thought to form amyloid-like fibers that anchor holdfast to the cell envelope. HfaB, a lipoprotein, is required for translocation of HfaA and HfaD to the cell surface. Deletion of the anchor proteins leads to a severe defect in adherence resulting from holdfast not properly attached to the cell and shed into the medium. This phenotype is greater in a Δ*hfaB* than a double Δ*hfaA hfaD* mutant, suggesting that HfaB has other functions besides the translocation of HfaA and HfaD. Here, we identify an additional HfaB-dependent holdfast anchoring protein, HfaE, which is predicted to be a secreted protein. HfaE is highly conserved among Caulobacterales species with no predicted function. In planktonic culture, *hfaE* mutants produce holdfasts and rosettes similar to wild type. However, holdfasts from *hfaE* mutants bind to the surface but are unable to anchor cells, similar to other anchor mutants. We showed that fluorescently-tagged HfaE co-localizes with holdfast, and HfaE forms an SDS-resistant high molecular weight species consistent with amyloid fiber formation. We propose that HfaE is a novel holdfast anchor protein, and that HfaE functions to link holdfast material to the cell envelope.

**IMPORTANCE:** For surface attachment and biofilm formation, bacteria produce adhesins that are composed of polysaccharides, proteins and DNA. Species in the Caulobacterales produce a specialized polar adhesin, holdfast, which is required for permanent attachment to surfaces. In this study, we evaluate the role of a newly identified holdfast anchor protein HfaE in holdfast anchoring to the cell surface in two different Caulobacterales with drastically different environments. We show that HfaE plays an important role in adhesion and biofilm formation in Caulobacterales. Our results provide insights into bacterial adhesins and how they interact with the cell envelope and surfaces.

## INTRODUCTION

Many bacteria spend their lives attached to or associated with surfaces, forming a complex community called a biofilm. Bacteria attach to surfaces using a variety of adhesins, which are mainly composed of polysaccharides, proteins, and DNA (1, 2). Most polysaccharide adhesins are synthesized in the cytoplasm and secreted by either an ABC-transporter dependent, a synthase-dependent, or a Wzx/Wzy-dependent translocation pathway, and can be associated with or anchored in the cell envelope (2–4). Protein adhesins are secreted via a Sec-dependent pathway involving an N-terminal leader peptide, and can be subsequently anchored to the cell surface by covalent linkage to the peptidoglycan in Gram-positive bacteria or the Gram-negative outer membrane, for example curli in *Escherichia coli* (5), protein F1 of *Streptococcus pyogenes* (6), or the fibronectin binding domain of *Staphylococcus aureus* (7). However, there are many other polysaccharide and protein adhesins whose anchoring systems are poorly understood.

Species of the order Caulobacterales use a polar adhesin called holdfast to adhere permanently to surfaces and form biofilms (2). The best characterized holdfast adhesin is from *C. crescentus,* a freshwater Caulobacterales*. Hirschia baltica*, a marine Caulobacterales, produces holdfast adhesin that binds better to surfaces in high ionic strength environments than the holdfast produced by *C. crescentus* (8). Holdfast is capable of binding to a variety of chemically distinct surfaces with impressive force (9–11). Although the complete composition of *C. crescentus* holdfast is unknown, it has been shown to contain *N*-acetylglucosamine (GlcNAc), glucose, 3-*O*-methylglucose, mannose, and xylose monosaccharides (12, 13), as well as proteins and DNA (14). *H. baltica* holdfast contains GlcNAc and galactose monosaccharides, and proteins (8).

The holdfast polysaccharide is produced via a mechanism similar to the Wzx/Wzy-dependent group I capsular polysaccharide synthesis pathway in *Escherichia coli* (15, 16). A putative glycosyltransferase HfsE initiates the synthesis of holdfast polysaccharide in the cytoplasm by transferring an activated sugar phosphate from uridine diphosphate (UDP) to an undecaprenyl-phosphate (Und-P) lipid carrier (16). Additional monosaccharide substituents are then added to form a repeat unit on the lipid carrier by three putative glycosyltransferases HfsG (16), HfsJ (17), and HfsL (18). Some of the sugar residues on the repeat units are enzymatically modified by a deacetylase, HfsH (19, 20), and a putative acetyltransferase, HfsK (21). The lipid carrier with the polysaccharide repeat unit is subsequently translocated across the inner membrane into the periplasm by the putative flippase HfsF (16). The repeat units are then predicted to be assembled into the mature polysaccharide in the periplasm by two polymerases, HfsC and HfsI (16). The assembled holdfast polysaccharide is then believed to be secreted through an export protein complex, composed of HfsA, HfsB, and HfsD (22–24).

Holdfast polysaccharides are anchored to the cell envelope by the action of the <u>h</u>oldfast <u>a</u>nchor (Hfa) proteins HfaA, HfaB, HfaD, and HfaE (18, 25–27). Deletion of anchor genes leads to holdfast shedding into the medium and a decrease in adhesion and subsequent biofilm formation (18, 25–27). HfaA, HfaB, and HfaD are colocalized at the tip of the stalk with the holdfast (26, 28). HfaA shares similarity with the curli monomer CsgA and forms a high molecular weight, SDS-resistant complex, a property of amyloid proteins (26). HfaA polymerization depends on HfaD, which shares limited sequence similarity to surface layer proteins and other adhesins (25, 26, 29). HfaB, a lipoprotein with sequence similarity to the curli secretin CsgG, is required for the stability of HfaA and HfaD and their localization to the cell surface (26). Loss of HfaA, HfaD, or both leads to partial shedding of holdfast into the medium, while loss of HfaB leads to severe holdfast shedding and abolishes biofilm formation (25, 26). It has been postulated that HfaB has a separate function in addition to facilitating the secretion of HfaA and HfaD, such as the formation of a complex with holdfast polysaccharides or secretion of other unidentified holdfast proteins (25, 26, 29). The mechanism by which holdfast anchor proteins and holdfast polysaccharides interact is poorly understood. However, deacetylation of holdfast polysaccharides is required for their tethering to the cell surface and may affect their association with the anchor proteins (20). We hypothesize that the amines generated as a result of deacetylation are important for the formation of bonds with anchor proteins, either covalently or electrostatically (20).

In this study, we characterize the HfaE protein in both *C. crescentus* and *H. baltica.* We show that HfaE is a secreted protein involved in holdfast anchoring. Deletion of *hfaE* in both *C. crescentus* and *H. baltica* reduces cell adhesion and biofilm formation; however, these mutants produce holdfasts and form rosettes similar to wild-type (WT) during planktonic growth. When cells are attached to a glass surface, even though the holdfasts produced by Δ*hfaE* mutants are able to adhere to the surface, the cells are easily washed away, indicating a holdfast anchoring defect. Finally, we show that HfaE colocalizes with holdfast and forms a high molecular weight, SDS resistant complex, a property of amyloids. We postulate that HfaE is a secreted holdfast protein that is required for holdfast to associate with the cell envelope, in addition to HfaA, HfaB, and HfaD.

## RESULTS

### HfaE is involved in cell adhesion

The mechanism by which holdfast polysaccharides are tethered to the cell envelope remains unknown. Three conserved *hfa* genes, *hfaA*, *hfaB*, and *hfaD* are found in the *hfa* operon in both *C. crescentus* and *H. baltica* (Fig. 1A). Shed holdfasts from double Δ*hfaA* Δ*hfaD* mutants contain peptides or proteins, suggesting that there are other unidentified holdfast proteins in addition to the known anchor proteins (8, 30). To identify uncharacterized holdfast genes, we explored genes adjacent to the *hfa* locus to test whether they may have a role in bacterial attachment or biofilm formation. *hbal_0649*, a predicted protein with unknown function, is immediately downstream of the *H. baltica hfa* locus (Fig. 1A). We performed reciprocal best-hit analysis and identified a homolog of *hbaI_0649* in the *C. crescentus* genome, *CC_2639 (CCNA_02722)*, which is eight genes downstream of the *C. crescentus hfa* locus (Fig. 1A). Hbal_0649 and CC_2639 are predicted to be secreted. Cells lacking CC_2639 were previously shown to be defective in surface attachment and holdfast anchoring in *C. crescentus* and the gene was named *hfaE* (18).

**Figure 1:**
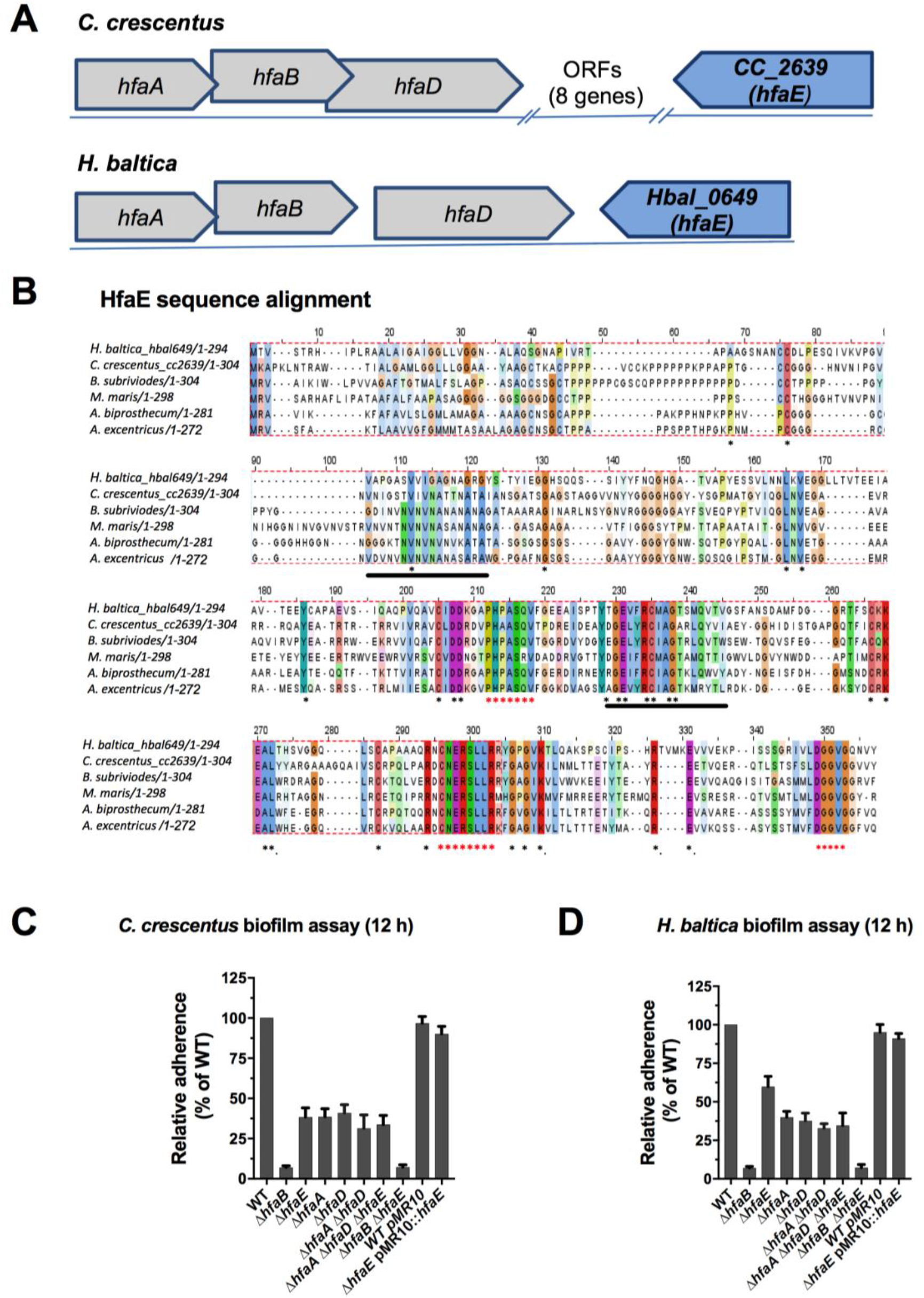
HfaE is involved in cell adhesion. **A.** Genomic organization of the holdfast anchor genes (*hfa)* in *C. crescentus* and *H. baltica.* Genes were identified using reciprocal best hit analysis with *C. crescentus* and *H. baltica* genomes. In the *C. crescentus* genome*, hfaE* is found outside the *hfa* locus, while in the *H. baltica* genome*, hfaE* is found downstream of *hfaD.* **B.** Multiple sequence alignment of HfaE from selected Caulobacterales. Residues are color coded based on their physicochemical properties using ClustalW (48). Consensus residues are indicated by an asterisk (*), and amyloid domains by black lines. **C-D.** Quantification of biofilm formation by the crystal violet assay after incubation for 12 h, expressed as a mean percent of WT crystal violet staining normalized to OD_600_. Error is expressed as the standard error of the mean of three independent biological replicates, each with four technical replicates.

We extended the reciprocal best-hit analysis to other Caulobacterales genomes and found that HfaE is highly conserved among the Caulobacterales. We identified *hfaE* gene in all fully sequenced Caulobacterales genomes (25 fully sequenced and annotated genomes) and in 204 out of 238 partially or drafted Caulobacterales genomes. Interestingly, HfaE and HfaA are two of the eleven proteins that are only found in Caulobacterales and not in any other sequenced genomes (31). Basic sequence analyses of HfaE proteins from different Caulobacterales species showed that HfaE has a highly conserved C-terminal region (half of the protein, Fig. 1B). HfaE from *C. crescentus* and *H. baltica* were 33% identical (ID) and 46% similar across their entire protein sequence. Although HfaE, HfaA, and HfaD all have similar overall sequence conservation between *C. crescentus* and *H. baltica* (∼ 33% ID), the C-terminus of HfaE is more conserved than average (∼43% ID), while the N-terminal region is more divergent (∼23% ID, 32). In particular, there are three C-terminal motifs that are highly conserved amongst all HfaE sequences analyzed: PHPASQV, CNERSLLR, and DGGVG (Fig. 1B, red asterisks). We hypothesized that these conserved motifs could be important for multimerization, protein-protein interactions, or protein-polysaccharide binding. We analyzed HfaE amino acid sequences using AGGRESCAN and PASTA2.0 to identify amino acid sequences important for promoting amyloid fiber formation (33, 34). We identified two common aggregation domains that are within the conserved regions of HfaE (Fig. 1B, black lines), suggesting HfaE might aggregate to form high molecular weight complexes similar to HfaA and HfaD.

We generated clean, in-frame deletions of *CC*_*2639 (hfaE_CC_)* and *hbal*_*0649 (hfaE_HB_)* in *C. crescentus* and *H. baltica* respectively, to determine whether they are involved in adhesion and/or biofilm formation. We found that deletion of *hfaE_CC_*in *C. crescentus* resulted in a 60% decrease in biofilm formation, similar to a Δ*hfaA* or Δ*hfaD* mutant (Fig. 1C). Deletion of *hfaE_HB_* in *H. baltica* led to a 40% reduction in biofilm formation compared to wildtype (WT), while a 60% reduction was observed for the *hfaA* and *hfaD* mutants (Fig. 1D). We complemented both the Δ*hfaE_CC_* and Δ*hfaE_HB_* mutants with their native copy of *hfaE* in *trans,* which restored biofilm formation to WT levels (Fig. 1C-D), confirming that HfaE is involved in biofilm formation in both *C. crescentus* and *H. baltica.* In *C. crescentus,* the triple Δ*hfaA* Δ*hfaD* Δ*hfaE* mutant phenocopies the double Δ*hfaA* Δ*hfaD* or each of the single mutants (Fig. 1C). In *H. baltica,* we observed that the triple Δ*hfaA* Δ*hfaD* Δ*hfaE* mutant phenocopies the double Δ*hfaA* Δ*hfaD,* Δ*hfaA,* and Δ*hfaD* mutants but not the Δ*hfaE* mutant, which has higher biofilm formation (Fig. 1D). These results suggest that HfaE does not contribute to biofilm formation independently of HfaA and HfaD and thus may function alongside or downstream of HfaA and HfaD. Importantly, the Δ*hfaA* Δ*hfaD* Δ*hfaE* triple mutant did not phenocopy the Δ*hfaB* mutant (Fig. 1C-D), suggesting that the additional hypothesized function of HfaB mentioned above is not just the secretion of HfaE. The Δ*hfaB* mutants were completely deficient in biofilm formation in both *C. crescentus* and *H. baltica*, which suggests that HfaE functions downstream of HfaB in holdfast anchoring, and may be involved in the same processes as HfaA and HfaD (Fig. 1C-D).

### HfaE contributes to holdfast anchoring

In order to study the role of HfaE in *C. crescentus* and *H. baltica* holdfast anchoring, we examined holdfasts using fluorescence microscopy with AF488 conjugated to the wheat germ agglutinin (WGA) lectin (AF488-WGA), which binds to GlcNAc moieties in the holdfast polysaccharide (12). In planktonic culture, *C. crescentus* and *H. baltica* WT cells formed organized clusters of cells attached together by their holdfast called rosettes. In addition, all holdfasts stained by AF488-WGA were attached to cells (Fig. 2A). The *C. crescentus* Δ*hfaE* and *H. baltica* Δ*hfaE* mutants were both labeled by AF488-WGA, indicating the presence of holdfast polysaccharide attached to cells (Fig. 2A). However, some holdfasts could also be detected away from any cell, suggesting that they were shed from cells in both mutant backgrounds (Fig. 2A, white arrows). Interestingly, when we quantified rosette formation in the Δ*hfaE* mutant, we observed that it was similar to WT (Fig. 2A, blue arrows and Fig. 2B), implying that holdfast anchoring in cells lacking HfaE was not severely impaired compared to the Δ*hfaA,* Δ*hfaD*, and Δ*hfaB* mutants which have a more reduced number of rosettes (Fig. 2B).

**Figure 2:**
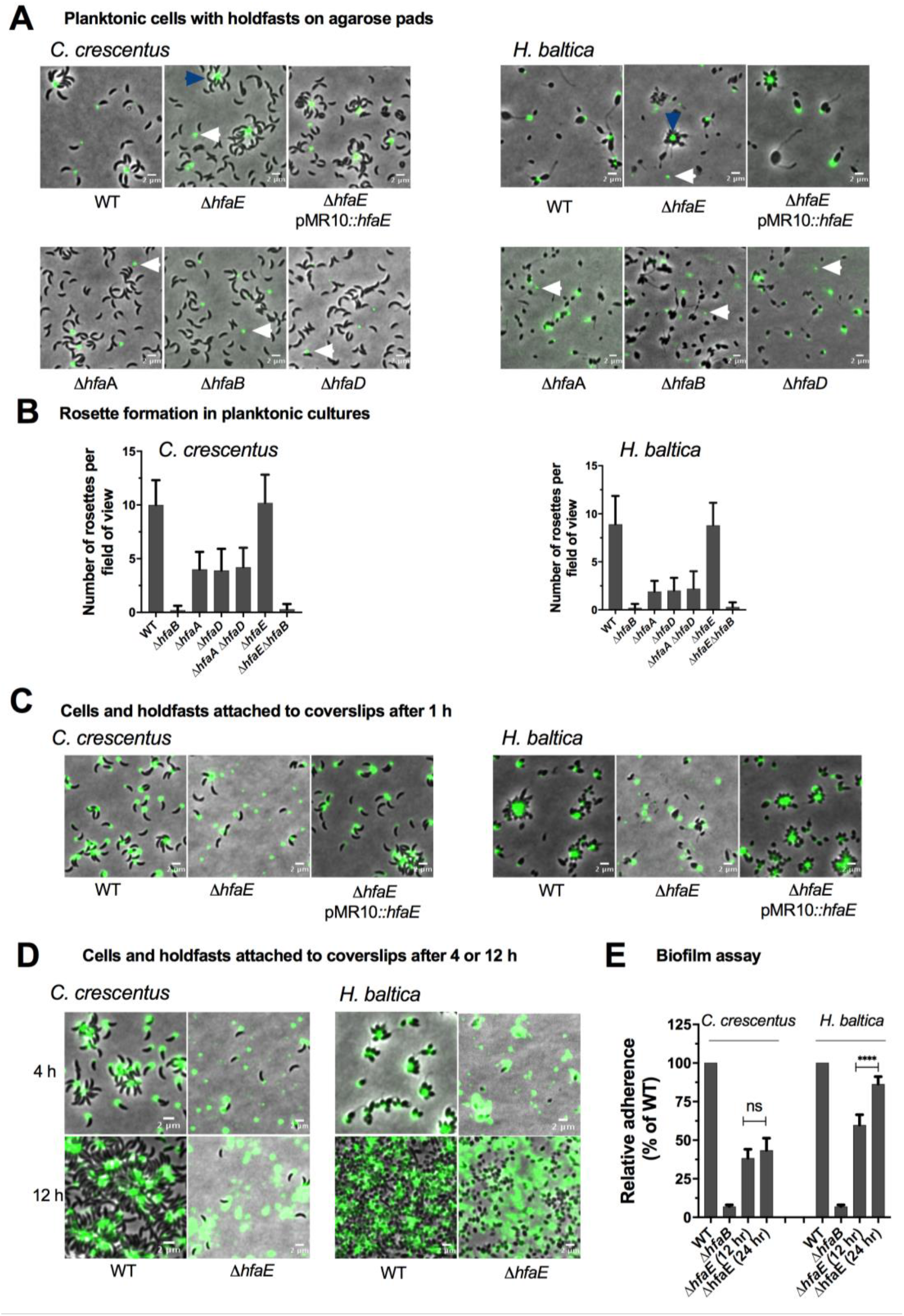
HfaE contributes to holdfast anchoring. **A.** Representative images showing merged phase and fluorescence channels of exponentially growing planktonic cultures of *C. crescentus* (left) and *H. baltica* (right) strains. Holdfast polysaccharides are labeled with AF488-WGA (green). White arrows indicate shed holdfast, while blue arrows indicate rosettes. **B**. Quantification of rosettes formed in *C. crescentus* (left) and *H. baltica* (right) strains grown to an OD_600_ of 0.8. Data are expressed as the mean number of rosettes formed. Error is represented as the standard error of the mean of two biological replicates with five technical replicates each. Scale bar, 2 µm**. C-D.** Representative images showing merged phase and fluorescence channels of *C. crescentus* and *H. baltica* strains bound to a glass slide. Holdfast is labeled with AF488-WGA (green). Exponentially growing cultures were incubated on the glass slides for 1 h (C), or 4 h to 12 h (D), and washed to remove unbound cells before AF488-WGA labeling. Scale bar, 2 µm**. E**. Quantification of biofilm formation by the crystal violet assay after incubation for 12 h and 24 h, expressed as a mean percentage of WT crystal violet staining normalized to OD_600_. Error is expressed as the standard error of the mean of three independent biological replicates, each with four technical replicates.

The reduction in biofilm formation in the Δ*hfaE* mutant was significant for both *C. crescentus* and *H. baltica* (Fig. 1C-D), however, we observed few shed holdfasts in the medium (Fig. 2A). This suggests that holdfasts from the Δ*hfaE* mutant may be loosely tethered to the cell surface. To test this hypothesis, we spotted exponentially growing cell cultures onto a glass coverslip and allowed cells to bind for 1 h. The coverslip was then washed to remove unbound cells and holdfast were labelled with AF488-WGA. As expected, WT cells were bound to coverslips via their holdfasts (Fig. 2C). However, for both *C. crescentus* and *H. baltica* Δ*hfaE* mutants, most cells were washed off the coverslip surface, leaving only their holdfasts attached (Fig. 2C). The strains complemented with *hfaE* in *trans* restored holdfast anchoring (Fig. 2C). These results indicate that HfaE plays an important role in holdfast anchoring for both *C. crescentus* and *H. baltica*.

We observed that both *C. crescentus* Δ*hfaE* and *H. baltica* Δ*hfaE* were strongly deficient in holdfast anchoring when cells were incubated with a surface for a short while (1 h, Fig. 2C), but *H. baltica* Δ*hfaE* showed increased cell adhesion after incubation with a surface for a longer period (4 h – 12 h, Fig. 2D). We hypothesized that over longer periods of time, the Δ*hfaE* cells are able to attach more efficiently to surfaces. To test this hypothesis, we quantified biofilm formation after 12 h and 24 h. We observed no significant increase in biofilm formed by *C. crescentus* Δ*hfaE* (Fig. 4E). However, *H. baltica* Δ*hfaE* showed an increase in biofilm formation to almost WT levels after 24 h (Fig. 4E). These results suggest that the role of HfaE varies between the two species, or that the contribution of HfaE to holdfast anchoring in both species differ due to differences in holdfast composition or holdfast anchoring mechanisms.

### Epistasis analysis of *hfaE* and other *hfa* genes

In the current model of holdfast anchoring in *C. crescentus,* HfaB forms a secretion pore through the outer membrane for the translocation of HfaA and HfaD to the cell surface, where they polymerize into high molecular weight fibers or complexes (26, 28). Recent studies on holdfast anchor mutants suggest that HfaB may have an additional function beyond the secretion of HfaA and HfaD (29). This was based on the observation that the loss of adhesion of a double *C. crescentus* Δ*hfaA* Δ*hfaD* mutant could be suppressed by mutations in sugar-nucleotide synthesis genes, whereas a *C. crescentus* Δ*hfaB* mutant could not be suppressed by similar mutations (29). The additional HfaB function is thought to be stabilization of the holdfast anchoring machinery, directly interacting with the polysaccharides or secretion of other unidentified anchor proteins.

We hypothesized that HfaB could be interacting with or transporting HfaE, thus deletion of *hfaE* in a Δ*hfaA* Δ*hfaD* background could increase holdfast shedding to levels observed in the Δ*hfaB* mutant. We monitored for the presence of holdfast in anchor mutants using AF488-WGA lectin. In planktonic culture for both *C. crescentus* and *H. baltica*, we observed shed holdfasts in all the anchor mutants (Fig. 3A). We quantified the number of shed holdfasts in planktonic cultures (Fig. 3B). In WT, it is very rare to observe shed holdfasts, while the Δ*hfaB* mutant has the highest shedding phenotype (Fig. 3B). We observed the lowest number of shed holdfasts in the Δ*hfaE* mutant compared to all the anchor mutants. The triple Δ*hfaA* Δ*hfaD* Δ*hfaE* mutant had a similar amount of shed holdfasts compared to the double Δ*hfaA* Δ*hfaD* mutant for both *C. crescentus* and *H. baltica*. These results suggest that HfaE may not be the only unidentified holdfast component that is secreted by HfaB, and further emphasize that HfaB may have additional unknown roles in holdfast anchoring.

**Figure 3:**
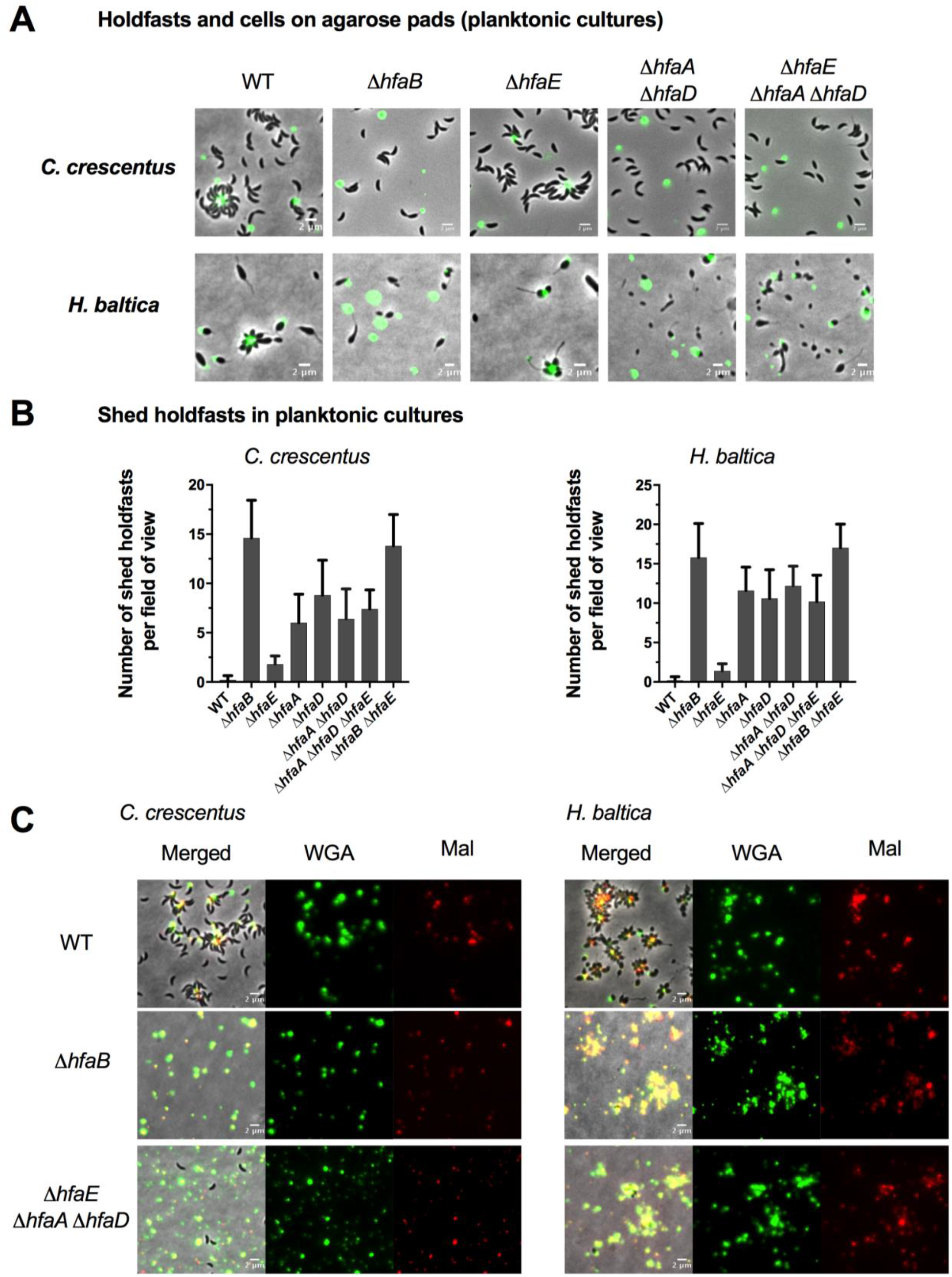
Epistasis analysis of *hfaE* and other *hfa* genes. **A.** Representative images showing merged phase and fluorescence channels of exponentially growing, planktonic *C. crescentus* (left) and *H. baltica* (right) strains on agarose pads. Holdfast polysaccharides are labeled with AF488-WGA (green). Scale bar, 2 µm**. B.** Quantification of shed holdfasts in *C. crescentus* (left) and *H. baltica* (right) strains grown to an OD_600_ of 0.8. Data are expressed as the mean number of shed holdfasts formed. Error is represented as the standard error of the mean of two biological replicates with five technical replicates each. **C.** Representative images showing merged phase and fluorescence channels of *C. crescentus* (left) and *H. baltica* (right) strains bound to a glass slide. Holdfast polysaccharides are labeled with AF488-WGA (GlcNAc, green) and holdfast thiols are labeled with the maleimide dye AF594-Mal (thiols, red). Exponential cultures were incubated on glass slides for 1 h, and washed to remove unbound cells before co-labelling with AF488-WGA and AF594-Mal. Scale bar, 2 µm.

Holdfasts from both *C. crescentus* and *H. baltica* are stained by maleimide, which indicates the presence of thiol components in the holdfast (8). To test whether the cysteines in HfaE are the thiol component of holdfasts, we allowed exponential-phase cultures to bind to coverslips for 1 h, washed to remove unbound cells, and co-labeled with both AF488-WGA (GlcNAc-specific, green) and the maleimide dye AF594-Mal (thiol-specific, red). As expected, the WT cells from *H. baltica* and *C. crescentus* were labeled with both AF488-WGA and AF594-Mal (Fig. 3C), indicating the presence of thiols in the holdfast. The shed holdfasts from the Δ*hfaB* mutants were also labelled with both AF488-WGA lectin and AF594-Mal (Fig. 3C). The single anchor mutants and the triple mutants Δ*hfaA* Δ*hfaD* Δ*hfaE* also showed similar labelling (Fig. 3C, Fig. S1). These results suggest that there are thiol components in the holdfast in addition to HfaA, HfaD, and HfaE cysteines, and that they are not secreted by HfaB.

### HfaE co-localizes with the holdfast and forms high molecular weight complexes

The holdfast anchor proteins HfaA, HfaD, and HfaB have been mainly studied in *C. crescentus* and they all colocalize with the holdfast polysaccharides (26). We decided to pursue our studies of HfaE in *C. crescentus* because its anchor proteins have been well characterized, and because *H. baltica* Δ*hfaE* has a less severe adhesion phenotype, complicating certain types of analyses. To determine the localization of HfaE in *C. crescentus*, HfaE was fused to mCherry at the C-terminus and fluorescent microscopy was used to examine HfaE::mCherry localization. We first confirmed that HfaE::mCherry is functional as it restores biofilm formation in *C. crescentus* Δ*hfaE* (Fig. S2). Next, an exponentially growing culture expressing HfaE::mCherry was labelled with AF488-WGA. We observed that HfaE::mCherry localized at the cell pole or at the tip of the stalk in stalked cells (Fig. 4A). In cells that have holdfasts (35 % of WT produce holdfasts under our growth conditions), HfaE::mCherry colocalized with holdfast (Fig. 4A). To test whether HfaE requires HfaA, HfaB, or HfaD to localize at the tip of the stalk, we generated in-frame deletions of *hfaA*, *hfaB* and *hfaD* in the *hfaE*::*mCherry* background. We added WGA to label holdfast and use fluorescent microscopy to observe HfaE::mCherry localization in these anchor mutants. In all the anchor mutants, HfaE::mCherry was mislocalized (Fig. 4A). These results indicate that HfaE requires HfaA, HfaD, and HfaB for correct localization at the tip of the stalk.

**Figure 4:**
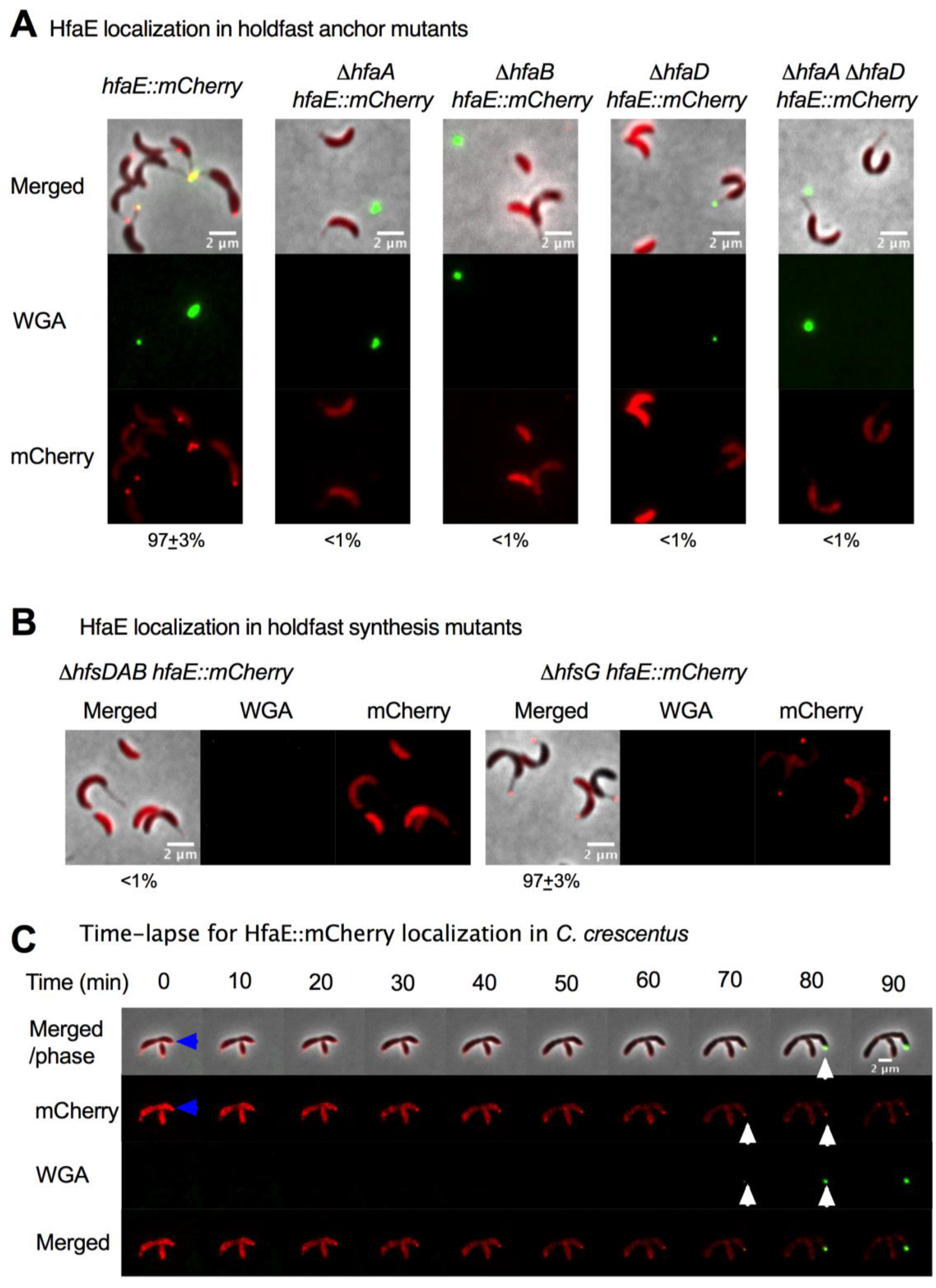
HfaE co-localizes with the holdfast in *C. crescentus*. **A-B.** Representative images showing merged phase and fluorescence channels of the indicated *C. crescentus* strains expressing HfaE::mCherry. Holdfasts were labeled with AF488-WGA (green), specific for GlcNAc in holdfast polysaccharides, and the mCherry channel (red) was used to visualize the localization of HfaE::mCherry. Exponential planktonic cultures were used to quantify the percentage of pre-divisional cells with HfaE::mCherry foci at the cell pole, which is indicated numerically at the bottom of each set of representative images. Data are expressed as the mean of two independent biological replicates, along with the standard error of the mean. A total of 5,000 cells were quantified per replicate using MicrobeJ. Scale bar, 2 µm**. C**. Time-lapse montages of *C. crescentus hfaE::mCherry* on soft agarose pads (0.8 % agarose). Exponential cultures were placed on soft agarose pads containing holdfast specific AF488-WGA (green) and covered with a glass coverslip. The blue arrow indicates a pre-divisional cell and white arrows indicate polar localization of HfaE::mCherry as well as the newly synthesized holdfast in the swarmer cell. Images were collected every 10 minutes for 6 hr. Scale bar, 2 µm.

HfaA, HfaD and HfaB localization at the tip of the stalk is independent of the presence of holdfast polysaccharides but depends on the holdfast secretion machinery (26). To test whether HfaE localization depends on the presence of holdfast polysaccharide or its secretion machinery, we generated a clean deletion of the holdfast export genes *hfsDAB* (no holdfast polysaccharide export protein complex and no holdfast produced) and a glycosyltransferase *hfsG* (presence of holdfast polysaccharide export protein complex but no holdfast polysaccharides) in the *hfaE::mCherry* background. We observed mislocalization of HfaE in Δ*hfsDAB* and correct localization in Δ*hfsG* (Fig. 4B), similarly to what has been observed with HfaA, HfaD, and HfaB localization (26). These results indicate that HfaE, like the other anchor proteins, requires the holdfast polysaccharide export complex but not the presence of the polysaccharide itself to localize at the tip of the stalk. In order to determine when HfaE localizes to the cell pole relative to when holdfast is synthesized, we performed time-lapse microscopy on soft agarose pads with AF488-WGA to label holdfast. We observed that HfaE::mCherry first localizes at cell pole in the pre-divisional cell (Fig. 4C, blue arrow). After cell division, HfaE::mCherry remains polarly localized in the swarmer cell where holdfast is synthesized (Fig. 4C, white arrows). This localization pattern is similar to what has been observed for the other anchor proteins (26).

HfaD and HfaA depend on each other for stability and localization (26). In order to test whether HfaA, HfaD, and/or HfaB localization depends on HfaE, we generated a clean deletion of *hfaE* in strains with C-terminally FLAG-tagged HfaA or HfaD (*hfaA::M2* and *hfaD::M2*). We performed immunofluorescence microscopy with an anti-FLAG antibody (IR-680), and observed that HfaA and HfaD correctly localized in Δ*hfaE* mutants (Fig. 5A). We also deleted *hfaE* in a strain expressing HfaB tagged with mCherry at its C-terminus (*hfaB::mCherry)*. We observed that HfaB localization is not affected in the Δ*hfaE* mutant (Fig. 5B). These results indicate that HfaA, HfaD, and HfaB localization do not depend on HfaE.

**Figure 5:**
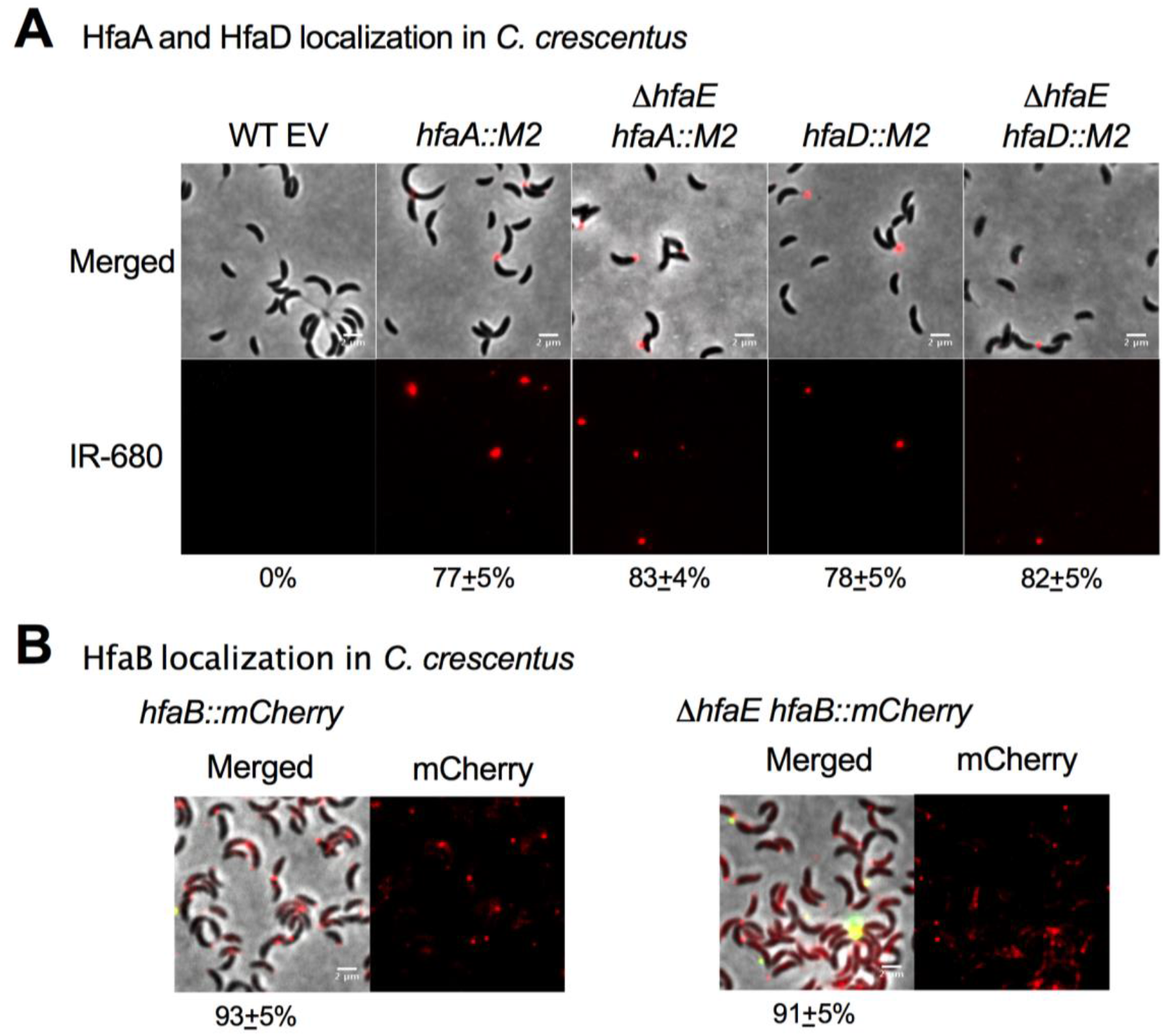
Localization of anchor proteins in *hfaE* mutants. **A.** Representative immunofluorescence images showing merged phase and fluorescence channels of *C. crescentus* strains expressing HfaA-M2 or HfaD-M2. Exponentially growing cells were fixed with formaldehyde. Anti-FLAG (M2) primary antibody and goat anti-rabbit secondary antibody conjugated to IRDye 680 red fluorescence was used to visualize localization of the anchor protein. **B**. Representative images showing phase and fluorescence channels of exponentially growing *C. crescentus hfaB::mCherry* strains. The mCherry fluorescence channel was used to visualize HfaB-mCherry (red). For panels **A-B**, exponential planktonic cultures were used to quantify the percentage of pre-divisional cells with fluorescent foci at the cell pole, which is indicated numerically at the bottom of each set of representative images. Data are expressed as the mean of two independent biological replicates, along with the standard error of the mean. A total of 1,000 cells were quantified per replicate using MicrobeJ. Scale bar, 2 µm.

HfaA and HfaD have been shown to form high molecular weight complexes that are resistant to heat and SDS denaturation, a characteristic of amyloid proteins (26). To determine if HfaE has similar properties, we performed western blot analysis of *C*. *crescentus hfaE*::*mCherry* whole cell lysates. We observed a high molecular weight species in the wells (Fig. 6A), similar to what was observed with HfaA and HfaD (26). To test whether the formation of this high molecular weight species depends on other anchor proteins, we deleted *hfaA, hfaB* and *hfaD* in the *C*. *crescentus hfaE::mCherry* background. We observed that HfaE multimerization was not affected in any of the anchor mutants (Fig. 6A, Fig.S3). These results suggest that HfaE assembles into a high molecular weight complex that is resistant to heat and SDS treatment, suggestive of an amyloid-forming protein (Fig. 1B). HfaA and HfaD depend on each other for stability and the formation of high molecular weight multimers (26). In order to test whether HfaE contributes to the stability of HfaA and HfaD, we performed western blots on strains with *hfaA::M2* and *hfaD::M2*. As expected, HfaA stability was affected in the *hfaD* mutant (Fig. 6B, upper panels). However, we observed that HfaA stability was not affected in the *hfaE* mutant (Fig. 6B, upper panels). We also observed similar results for HfaD (Fig. 6B, lower panels). These results indicate that HfaA and HfaD do not depend on HfaE for multimerization and stability.

**Figure 6:**
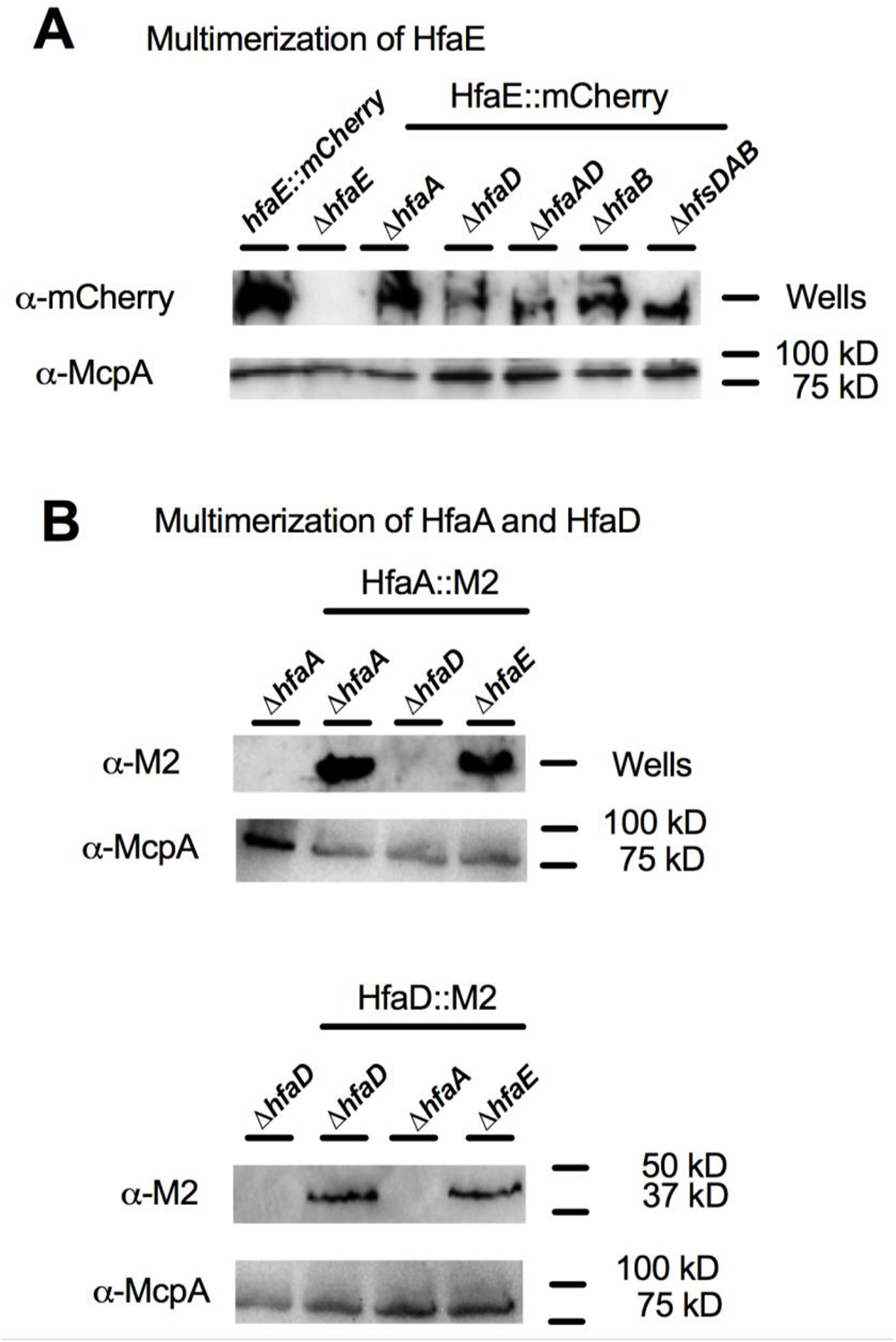
HfaE forms high molecular weight complexes. **A-B**. Western blots of whole cell lysates of *C. crescentus hfaE::mCherry* (A), *C. crescentus hfaA::FLAG* and *C. crescentus hfaD::FLAG* (B) using anti-FLAG tag (α-M2) antibody with goat anti-rabbit HRP conjugated secondary antibody. Exponentially growing cells grown to an OD_600_ of 0.6-0.8 were normalized to the equivalent of 1 mL of culture at an OD_600_ of 1.0, and 10 ul of the cell lysate was loaded into each lane. Anti-McpA was used as a loading control.

### Mutations in sugar-nucleotide synthesis pathways suppress the *hfaE* mutation

Little is known about how the holdfast anchor complex interacts with holdfast polysaccharides. Deacetylation of holdfast polysaccharides is required for strong interactions between the holdfast polysaccharides, anchor complex, and the thiol-containing component(s) of the holdfast (19, 20). Mutations in sugar-nucleotide synthesis genes have been shown to suppress holdfast shedding in *hfaA* and *hfaD* mutants, but not a *hfaB* mutant (29). These mutations are hypothesized to alter lipopolysaccharide (LPS) properties on the cell envelope, permitting an alternative mechanism for holdfast polysaccharide to interact with the cell envelope (18, 29). To test whether similar mutations can suppress the holdfast shedding phenotype of the *hfaE* mutant, we generated clean deletions of *wbqV* (dehydratase that catalyzes the conversion pf UDP-GlcNAC to UDP-Qui2NAc) and *rfbB* (dTDP-glucose 4,6-dehydratase that converts dTDP-D-glucose to dTDP-6-deoxy-D-glucose) in the *C. crescentus ΔhfaE* background. We allowed cells to bind to a glass surface for 1 h, added AF488-WGA, and washed away unattached cells. We observed that both *wbqV* and *rfbB* mutations suppressed holdfast shedding in the *hfaE* mutant (Fig. 7A). We observed similar restoration of adhesion and biofilm formation to WT levels in both Δ*wbqV* Δ*hfaE* and Δ*rfbB* Δ*hfaE* mutants (Fig. 7B).

**Figure 7:**
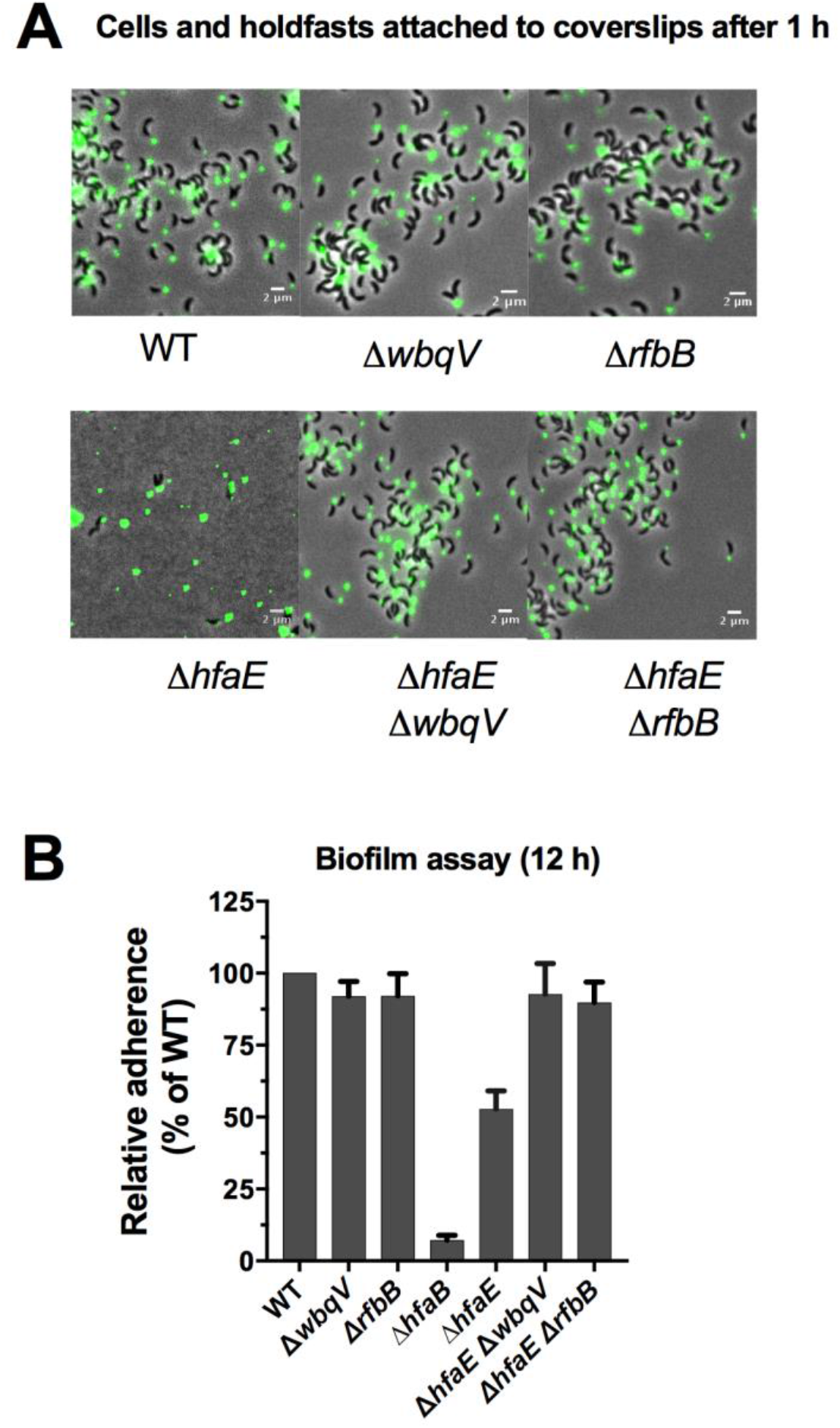
Mutations in sugar-nucleotide synthesis genes suppress the *hfaE* mutation. **A.** Representative images showing merged phase and fluorescence channels of exponentially growing, planktonic *C. crescentus* strains. Exponentially growing cultures were incubated on glass slides for 1 h, and washed to remove unbound cells before labelling with AF488-WGA (GlcNAc, green). Scale bar, 2 µm**. B.** Quantification of biofilm formation by the crystal violet assay after incubation for 12 h, expressed as a mean percent of WT crystal violet staining normalized to OD_600_. Error is expressed as the standard error of the mean of three independent biological replicates, each with four technical replicates.

## DISCUSSION

Reversible attachment of bacteria to surfaces is often facilitated by pili and flagella, while permanent adhesion is mediated by synthesized adhesins composed of exopolysaccharides and/or proteins (1, 35). Members of the Caulobacterales use a polar adhesin composed of polysaccharides and proteins called the holdfast to permanently attach to surfaces and form biofilms (1, 2, 8, 12, 36). Holdfast polysaccharides are tethered to the cell surface by the anchor proteins HfaA, HfaB, and HfaD (25, 26). HfaB is predicted to be an outer membrane localized pore-forming lipoprotein which facilitates secretion of HfaA and HfaD (22, 26). Because the holdfast anchoring defect of a *hfaB* mutant is much more severe than that of a double Δ*hfaA* Δ*hfaD* mutant (25, 26), we hypothesized that there were still unidentified holdfast anchor genes. By analyzing the synteny of holdfast anchor genes, we identified *CC_2639* from *C. crescentus* and *hbal*_*0649* from *H. baltica* as putative additional anchor genes. While this work was in progress, Hershey *et al.* identified *CC_2639 (hfaE)* in *C. crescentus* as a gene whose deletion causes holdfast shedding (18).

HfaE has a signal sequence that is predicted to target it to the periplasm or outer membrane. Deletion of *hfaE* in *C. crescentus* decreases biofilm formation to half that of WT, similar to deletion of *hfaA* or *hfaD*. In *H. baltica*, deletion of *hfaE* decreases biofilm formation, but not to the same levels as the Δ*hfaA* or Δ*hfaD* mutants. These results indicate that HfaE is involved in bacterial adhesion, although its contribution appears to differ between species. This could be due to differences in anchoring mechanisms, since holdfasts in *H. baltica* are anchored to the cell pole and cover a larger area compared to *C. crescentus* holdfasts, which are anchored at the tip of the thin stalk. Alternatively, the observed differences could be due to the differences in holdfast composition and structure. *H. baltica* holdfasts contain galactose monosaccharides, which are absent in *C. crescentus* (8).

We showed that cells lacking HfaE shed few holdfasts into the medium during planktonic growth, and that cells bearing a holdfast form rosettes similarly to WT. However, when cells are attached to a glass coverslip surface, most Δ*hfaE* cells detached from the coverslip after washing, leaving only their holdfasts. Over longer periods of growth, *H. baltica* Δ*hfaE* was able to increase the proportion of cells attached to coverslips, while *C. crescentus* Δ*hfaE* remained poorly attached. These results imply that the role of HfaE is to strengthen the association between other holdfast matrix components and the cell envelope, and that this requirement is variable between these species. HfaE::mCherry colocalizes with holdfast in *C. crescentus*, similar to HfaA and HfaD (26), suggesting that it may be part of the anchor complex. Deletion of *hfaB* in both *C. crescentus* and *H. baltica* leads to severe holdfast shedding and abolishes biofilm formation (25, 26). It was intriguing that a Δ*hfaA* Δ*hfaD* Δ*hfaE* triple mutant did not phenocopy a Δ*hfaB* mutant, suggesting that there are likely other unidentified anchor proteins or that HfaB interacts directly with holdfast polysaccharides. Holdfast has been shown to contain thiols other than the cysteines of holdfast anchor proteins HfaA and HfaD based on labelling with a thiol-reactive maleimide dye (8, 9). We hypothesized that HfaE could be the main thiol-containing component of holdfast since it has 10 cysteine residues, however our results showed that this is not the case.

Using western blot analysis, we showed that HfaE forms a high molecular weight complex that is resistant to heat and SDS denaturation, similar to HfaA, HfaD, and other amyloid-forming proteins (26). HfaA and HfaD depend on each other for stability and multimerization (26). Our results show that HfaE depends on HfaA, HfaB, and HfaD for localization to the cell pole, however, HfaE forms a high molecular weight complex without these anchor proteins. This suggests that either HfaE self-polymerizes, or that there is an unidentified protein involved in HfaE assembly, similarly to the way HfaA and HfaD depend on each other for stability and polymerization.

How holdfast polysaccharides are attached to the cell envelope via the anchor complex is still unknown. Polysaccharide deacetylase mutants shed holdfast similarly to anchor mutants, suggesting that deacetylation of holdfast polysaccharides is required for strong interactions between the holdfast polysaccharides and the anchor complex (19, 20). Mutations in the sugar-nucleotide synthesis genes, *wbqV* and *rfbB,* suppress holdfast shedding and restore biofilm formation in the *hfaE* mutant, similar to what has been observed in the *hfaA* and *hfaD* mutants (29). These mutations in the sugar-nucleotide synthesis pathway are hypothesized to (1) alter LPS properties on the cell envelope, permitting an alternative mechanism for holdfast polysaccharide to interact with the cell envelope or (2) alter holdfast polysaccharide composition and increases its interaction with some other components of the cell envelope (18, 29). However, these sugar-nucleotide biosynthetic mutations do not suppress the *hfaB* mutation (29). This result suggests that HfaE is a part of the anchor complex along with HfaA and HfaD, and that HfaB is directly interacting with the polysaccharides or that there are other unidentified anchor proteins that play a role in suppression of the *hfaA, hfaD,* and *hfaE* mutations.

Even though HfaE is functionally similar to HfaA and HfaD, there are some differences in the phenotypes of mutants in each of these genes which suggest that they play slightly different roles, despite all co-localizing with the holdfast polysaccharides and forming multimeric complexes. We propose that HfaE strengthens the anchor complex, and that in the absence of HfaE, the anchor complex (HfaA-HfaB-HfaD) is still able to partially tether the holdfast polysaccharides to the cell envelope, but that this anchoring is unable to withstand shear forces. Whether these proteins are all part of the same anchoring complex, and how they contribute to holdfast anchoring individually and collectively, will be the focus of future studies.

## MATERIALS AND METHODS

### Bacterial strains and growth conditions

The bacterial strains used in this study are listed in Table S1. *H. baltica* strains were grown in marine medium (Difco^TM^ Marine Broth/Agar 2216). *C. crescentus* was grown in PYE medium. Both *H. baltica* and *C. crescentus* strains were grown at 30 °C. When appropriate, antibiotics were added at the following concentrations: kanamycin, 5 µg/ml in liquid medium and 20 µg/ml in agar plates; streptomycin, 5 µg/ml in liquid medium and in plates. *E. coli* strains were grown in LB medium (37) at 37 °C and supplemented with 30 µg/ml of kanamycin in liquid medium or 25 µg/ml in agar plates; 100 µg/ml of spectinomycin in liquid medium or 50 µg/ml in agar plates; and 12 µg/ml of chloramphenicol in liquid medium and in agar plates.

### Strain construction

All the plasmids and primers used in this study are listed in Table S1 and S2, respectively. In-frame deletion mutants were generated by double homologous recombination as previously described (38) using suicide plasmids transformed into the *C. crescentus* or *H. baltica* host strains by electroporation (39). Briefly, genomic DNA was used as the template to PCR-amplify 500 bp fragments upstream and downstream of the gene to be deleted. The primers were designed with 25 bp of overlapping sequence for isothermal assembly (40) into pNPTS139 plasmid digested using EcoRV-HF (New England Biolabs, Ispswich, MA) using the NEBuilder HiFi Assembly Master mix (NEB). Assembled pNPTS139-based constructs were transformed into *α-*select *E. coli* for screening and sequence confirmation before introduction into the host *C. crescentus* or *H. baltica* strains by electroporation. Introduction of the desired mutation onto the *C. crescentus* or *H. baltica* genome was verified by sequencing.

For gene complementation, pMR10 was digested with EcoRV-HF and 500 bp upstream of *hfaE* containing the promoter elements, and the *hfaE* gene itself, were ligated into pMR10 using NEBuilder HiFi Master mix (NEB). The pMR10-based constructs were transformed into *α-*select *E. coli* for screening and sequence confirmation before introduction into either *C. crescentus* or *H. baltica* by electroporation, followed by selection for kanamycin resistance.

For generation of the HfaE::mCherry construct, pCHYC-1 and a 500 bp ‘*hfaE* C-terminal PCR product were digested with HindIII-HF and KpnI (NEB) and ligated with T4 DNA ligase (NEB). The pCHYC-1-*hfaE*::mCherry construct was transformed into *α-*select *E. coli* for screening and sequence confirmation before conjugation into *C. crescentus* using *E. coli* SM10, a S17-1 derivative. Transconjugants were identified via streptomycin resistance.

### Holdfast labeling using fluorescent lectins

Holdfast labeling with AF488-WGA (ThermoFisher, Waltham, MA) was performed as previously described (8) with the following modifications. Overnight cultures were diluted in fresh medium to an OD_600_ of 0.2 and incubated for 4 h to an OD_600_ of 0.6 – 0.8. AF488-WGA was added to 100 µl of the resultant exponential culture to a final concentration of 0.5 µg/ml and incubated at room temperature for 5 min. Five microliters of the labeled culture was then spotted onto a glass cover slide, covered with a 1.5 % (w/v) agarose (Sigma-Aldrich) pad in water, and visualized by epifluorescence microscopy. Imaging was performed using an inverted Nikon Ti-E microscope with a Plan Apo 60X objective, a GFP/DsRed filter cube, an Andor iXon3 DU885 EM CCD camera, and Nikon NIS Elements imaging software with 200 ms exposure time. Images were processed in ImageJ (41).

### Biofilm formation assays

Biofilm formation assays were performed as previously described (8) with the following modifications. For short-term binding assays, exponential cultures (OD_600_ of 0.6 – 0.8) were diluted to an OD_600_ of 0.4 in fresh marine broth (*H. baltica*) or PYE (*C. crescentus*), added to a 24-well plate (1 mL per well), and incubated with shaking (100 rpm) at room temperature for 4 h. For biofilm assays, overnight cultures were diluted to an OD_600_ of 0.1, added to a 24-well plate (1 mL per well), and incubated at room temperature for 12 h. with shaking (100 rpm). In both set-ups, OD_600_ was measured before the wells were rinsed with distilled H_2_O to remove non-attached bacteria, stained using 0.1% crystal violet (CV), and rinsed again with dH_2_O to remove excess CV. The CV was dissolved with 10% (v/v) acetic acid and quantified by measuring the absorbance at 600 nm (A_600_). Biofilm formation was normalized to A_600_ / OD_600_ and expressed as a percentage of WT.

### Visualization of holdfasts attached to glass coverslips

Visualization of holdfast binding to coverslips was performed as described previously (8) with the following modifications. *H. baltica* and *C. crescentus* strains grown to exponential phase (OD_600_ of 0.4 – 0.6) were incubated on washed glass coverslips at room temperature in a saturated humidity chamber for 4 – 8 h. After incubation, the slides were rinsed with dH_2_O to remove unbound cells and holdfasts were labelled using 50 µL of AF488-WGA at a concentration of 0.5 µg/ml for 5 min at room temperature. Excess lectin was washed off and the slides were topped with a glass coverslip. Holdfasts were imaged by epifluorescence microscopy using an inverted Nikon Ti-E microscope with a Plan Apo 60X objective, a GFP/DsRed filter cube, an Andor iXon3 DU885 EM CCD camera, and Nikon NIS Elements imaging software with 200 ms exposure time. Images were processed in ImageJ (41). Dual labelling with HfaE::mCherry and AF488-WGA was performed as previously described in Hardy et al 2010.

### Holdfast labeling using fluorescent maleimide

Alexa Flour conjugated Maleimide C_5_ (AF594-mal, ThermoFisher Scientific) was added to 100 µl of exponential culture to a final concentration of 0.5 µg/ml and incubated at room temperature for 5 min. A 5 µl aliquot of the labeled culture was spotted onto a glass coverslide, covered with a 1.5 % (w/v) agarose (Sigma-Aldrich) pad in water, and visualized by epifluorescence microscopy. Imaging was performed using an inverted Nikon Ti-E microscope with a Plan Apo 60X objective, a GFP/DsRed filter cube, an Andor iXon3 DU885 EM CCD camera, and Nikon NIS Elements imaging software with 200 ms exposure time. Images were processed in ImageJ (41).

### Holdfast synthesis and HfaE localization by time-lapse microscopy on soft agarose pads

*C. crescentus* holdfast synthesis was observed in live cells on agarose pads by time-lapse microscopy as described previously (8) with some modifications. A 1 µl aliquot of exponential-phase cells (OD_600_ of 0.4 – 0.8) was placed on top of a 0.8% agarose pad in PYE with 0.5 µg/ml of AF488-WGA. The pad was overlaid with a coverslip and sealed with VALAP (Vaseline, lanolin and paraffin wax). Time-lapse microscopy images were taken every 10 min for 6 h using an inverted Nikon Ti-E microscope and a Plan Apo 60X objective, a GFP/DsRed filter cube, and an Andor iXon3 DU885 EM CCD camera. Time-lapse movies were processed using ImageJ (41).

### Sample preparation, SDS-PAGE, and Western blot analysis

Cell lysates were prepared from exponentially growing cultures (OD_600_ 0.6-0.8) as previously described in (26), with the following modifications. The equivalent of 1.0 ml of culture at OD_600_ 0.6-0.8 was centrifuged at 16,000 × *g* for 5 min at 4 °C. The supernatant was removed, and cell pellets were resuspended in 50 µl of 10mM Tris, pH 8.0. 50 µl of 2x SDS sample buffer was then added to the cell suspension. Samples were boiled for 5 min at 100 °C before separation on a 12% (w/v) polyacrylamide gel and transfer to a nitrocellulose membrane. Membranes were blocked for 30 min in 5% (w/v) non-fat dry milk in TBST (20 mM Tris, pH 8, 137 mM NaCl, and 0.05% (w/v) Tween 20), and incubated at 4 °C overnight with primary antibodies. Anti-FLAG tag or anti-mCherry primary antibodies were used at a concentration of 1:1,250 (Sigma, St. Louis, MO). Then, a 1:20,000 dilution of HRP-conjugated goat anti-rabbit secondary antibody (Bio-Rad, Hercules, CA), was incubated with the membranes at room temperature for 1 h. Membranes were developed with SuperSignal West Dura Substrate (Thermo Scientific, Rockford, IL) and imaged with a Bio-Rad Chemidoc MP imager (Bio-Rad).

### Immunofluorescence microscopy

Immunofluorescence microscopy was performed as previously described (42) for unpermeabilized cells with some modification. Briefly, cells were grown to an OD_600_ of 0.3–0.8 and fixed in 2.5% formaldehyde (Ted Pella, Redding, CA) for 15 min. Cells were spun at 8,000 *g* for 5 min at 4°C and washed thrice with phosphate-buffered saline (PBS), pH 7.2 and resuspended in 0.5% (w/v) blocking reagent from Roche Molecular Biochemicals in 1× PBS for 30 min at 37°C. Primary antibody (anti-FLAG) was added (1:100 dilution) and incubated for 2 h. at 37°C. Cells were washed three times in 1× PBS. Secondary antibody [fluorescein isothiocyanate (FITC)-conjugated goat anti-rabbit; Jackson Immunological Research, West Grove, PA] was added (1:100) in blocking buffer and incubated for 1 h. at 37°C. Cells were washed three times in PBS−0.05% Tween 20. Imaging was performed using an inverted Nikon Ti-E microscope with a Plan Apo 60X objective, a GFP/DsRed filter cube, an Andor iXon3 DU885 EM CCD camera, and Nikon NIS Elements imaging software with 200 ms exposure time. Images were processed in ImageJ (41).

## ACKNOWLEDGEMENTS

We thank the members of the Brun laboratory for the discussion and providing critical comments on the manuscript. This study was supported by grant R35GM122556 from the National Institutes of Health to YVB. YVB is supported by a Canada 150 Research Chair in Bacterial Cell Biology from the Canadian Institutes of Health Research.

---

**TABLE S1:** Bacterial strains and plasmids.

| Strain or Plasmid | Description and/or genotype | Reference / source |
| --- | --- | --- |
| *E. coli* |  |  |
| $\alpha$ select | F<sup>-</sup>*deoR endA1 relA1 gyrA96 hsdR17(r<sub>K</sub><sup>-</sup>m<sub>K</sub><sup>+</sup>) supE44 thi-1 $\Delta$(lacZYA-argFV169) Φ80δlacZΔM15 $\lambda$* | Bioline |
| S17-1 | *E. coli* 294::RP4-2(Tc::Mu)(Km::Tn7) | (43) |
| SM10 | *thi-1 thr leu tonA supE recA*::RP4-2 Tc::Mu, KmR | (43) |
| *C. crescentus* |  |  |
| YB135 | Wild-type strain CB15 | (44) |
| YB4250 | CB15 Δ*hfaA* | (26) |
| YB4251 | CB15 Δ*hfaB* | (26) |
| YB4252 | CB15 Δ*hfaD* | (26) |
| YB4288 | CB15 Δ*hfaA* Δ*hfaD* | (26) |
| YB | CB15 pJU142 | (26) |
| YB5618 | CB15 Δ*hfaA* pJU142::*hfaAM2* | (26) |
| YB5620 | CB15 Δ*hfaD* pJU142::*hfaDM2* | (26) |
| YB5637 | CB15 Δ*hfaB* pCHYC-1::*hfaABmCherry* | (26) |
| YB188 | CB15 Δ*hfaE* | This study |
| YB197 | CB15 Δ*hfaE* pMR10:*hfaE* | This study |
| YB204 | CB15 Δ*hfaA* Δ*hfaD* Δ*hfaE* | This study |
| YB8734 | CB15 *hfaE*::pCHYC-1::*hfaEmCherry* | This study |
| YB8735 | CB15 *hfaE*::pJM21::*hfaEM2* | This study |
| YB9557 | CB15 Δ*hfaE* Δ*hfaA* pJU142::*hfaAM2* | This study |
| YB9558 | CB15 Δ*hfaE* Δ*hfaD* pJU142::*hfaDM2* | This study |
| YB9614 | CB15 Δ*hfaA* *hfaE*::pJM21::*hfaEM2* | This study |
| YB9615 | CB15 Δ*hfaD* *hfaE*::pJM21::*hfaEM2* | This study |
| YB9617 | CB15 Δ*hfaA* Δ*hfaD* *hfaE*::pJM21::*hfaEM2* | This study |
| YB9618 | CB15 Δ*hfaB* *hfaE*::pJM21::*hfaEM2* | This study |
| YB9619 | CB15 Δ*hfaE* Δ*hfaB* pCHYC-1::*hfaBmCherry* | This study |
| YB9627 | CB15 Δ*hfaE* Δ*rfbB* | This study |
| YB9628 | CB15 Δ*hfaE* Δ*wbqV* | This study |
| *H. baltica* |  |  |
| YB5842 | IFAM 1418 Wild-type strain | (45) |
| YB8404 | YB5842 Δ*hfaB* | (8) |
| YB8425 | YB5842 Δ*hfaD* | (8) |
| YB176 | YB5842 Δ*hfaE* | This study |
| YB191 | YB5842 Δ*hfaE* pMR10:*hfaE* | This study |
| YB186 | YB5842 Δ*hfaA* | (8) |
| YB208 | YB5842 Δ*hfaA* Δ*hfaD* | (8) |
| YB209 | YB5842 Δ*hfaA* Δ*hfaD* Δ*hfaE* | This study |
| **Plasmids** |  |  |
| pMR10 | Mini-RK2 cloning vector; RK2 replication and stabilization functions, KmR | R. Roberts and C. Mohr |
| pCHYC-1 | integrating plasmid with C-terminal mCherry fusion, StrR/SpR | (46) |
| pJM21 | integrating plasmid with C-terminal M2tag; KmR | (47) |
| pLVC9 | conjugation helper plasmid carrying a ColEI mob, CmR | G. Warren, unpublished |
| pMR10:*hfaE*_CC | pMR10 containing *hfaE* gene from *C. crescentus* for complementation | This study |
| pMR10:*hfaE*_HB | pMR10 containing *hfaE* gene from *H. baltica* for complementation | This study |
| pCHYC-1::*hfaE* | integrating plasmid containing a 500 bp C-terminal *hfaE* fragment with a C-terminal mCherry tag | This study |
| pJM21::*hfaEM2* | integrating plasmid containing a 500 bp C-terminal *hfaE* fragment with a C-terminal M2 tag | This study |
| pNPTS139 | pLitmus 39 derivative, *oriT*, *sacB*, Kan<sup>r</sup> | M.R.K. Alley, unpublished |
| pNPTS139Δ*hfaE*_CC | pNPTS139 containing 500 bp fragments upstream and downstream of *C. crescentus* *hfaE* | This study |
| pNPTS139Δ*hfaE*_HB | pNPTS139 containing 500 bp fragments upstream and downstream of *H. baltica hfaE* | This study |
| pNPTS139ΔrfbB | pNPTS139 containing 500 bp fragments upstream and downstream of *C. crescentus rfbB* | (29) |
| pNPTS139ΔwbqV | pNPTS139 containing 500 bp fragments upstream and downstream of *C. crescentus rfbB* | (29) |

**TABLE S2:**
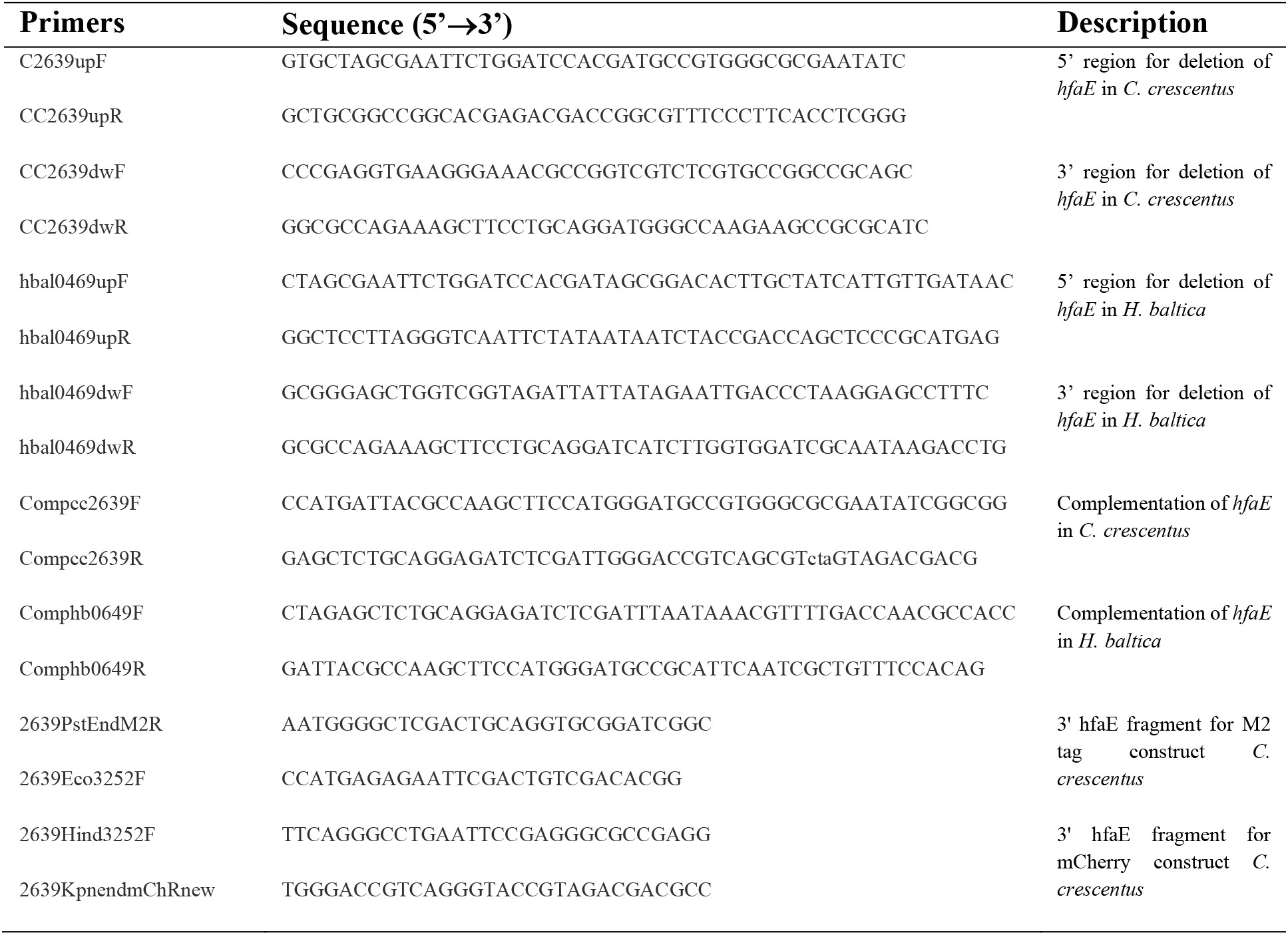
Primers.

**Figure S1:**
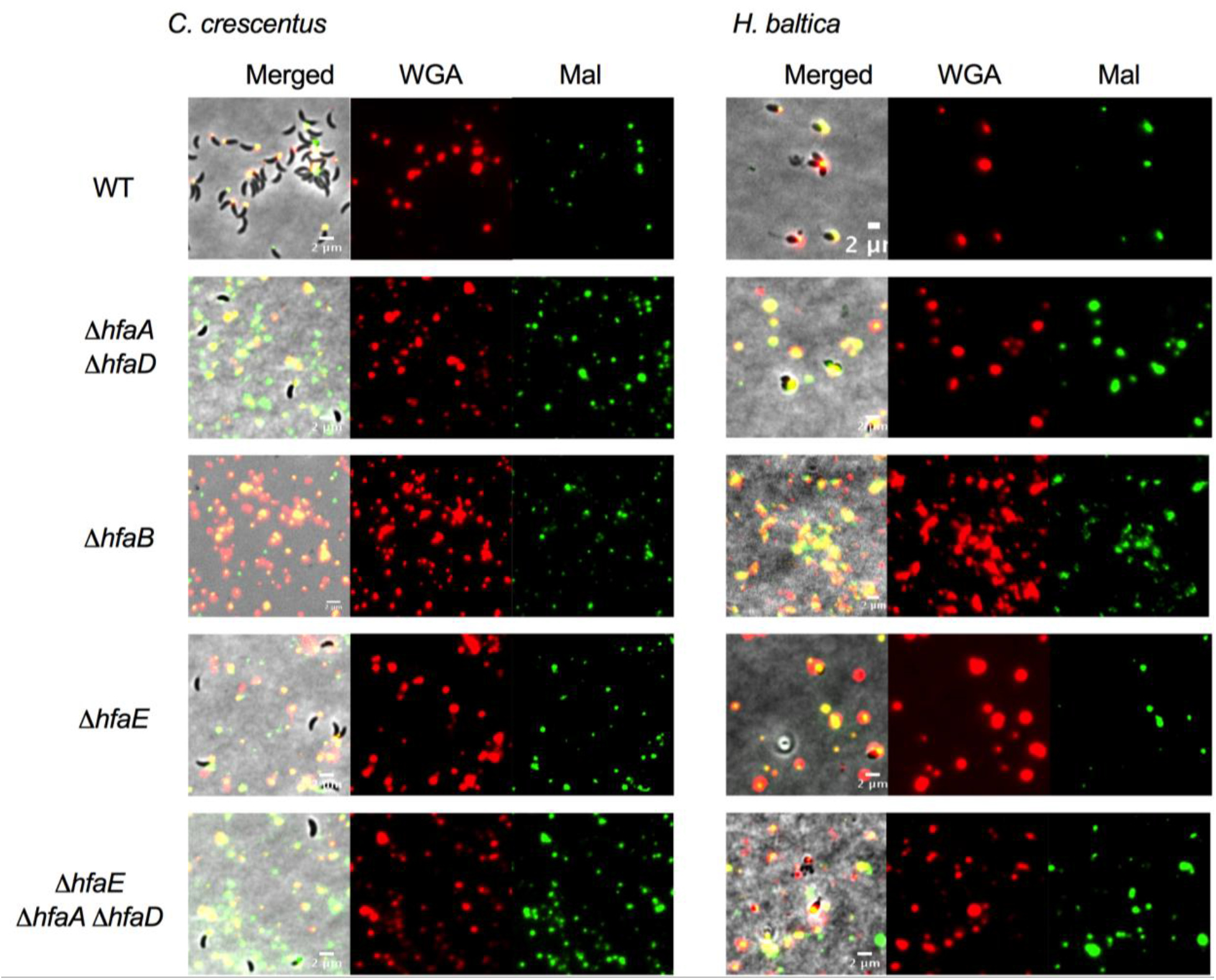
Thiol labeling in holdfasts, related to figure 3. Representative images showing merged phase and fluorescence channels of exponentially growing, planktonic *C. crescentus* (left) and *H. baltica* (right) strains. Holdfast polysaccharides are labeled with AF488-WGA (GlcNAc, green) and holdfast thiols are labeled with the maleimide dye AF594-Mal (thiols, red). For surface attachment, exponential cultures were incubated on glass slides for 1 h, and washed to remove unbound cells before co-labelling with AF488-WGA and AF594-Mal. Scale bar, 2 µm.

**Figure S2:**
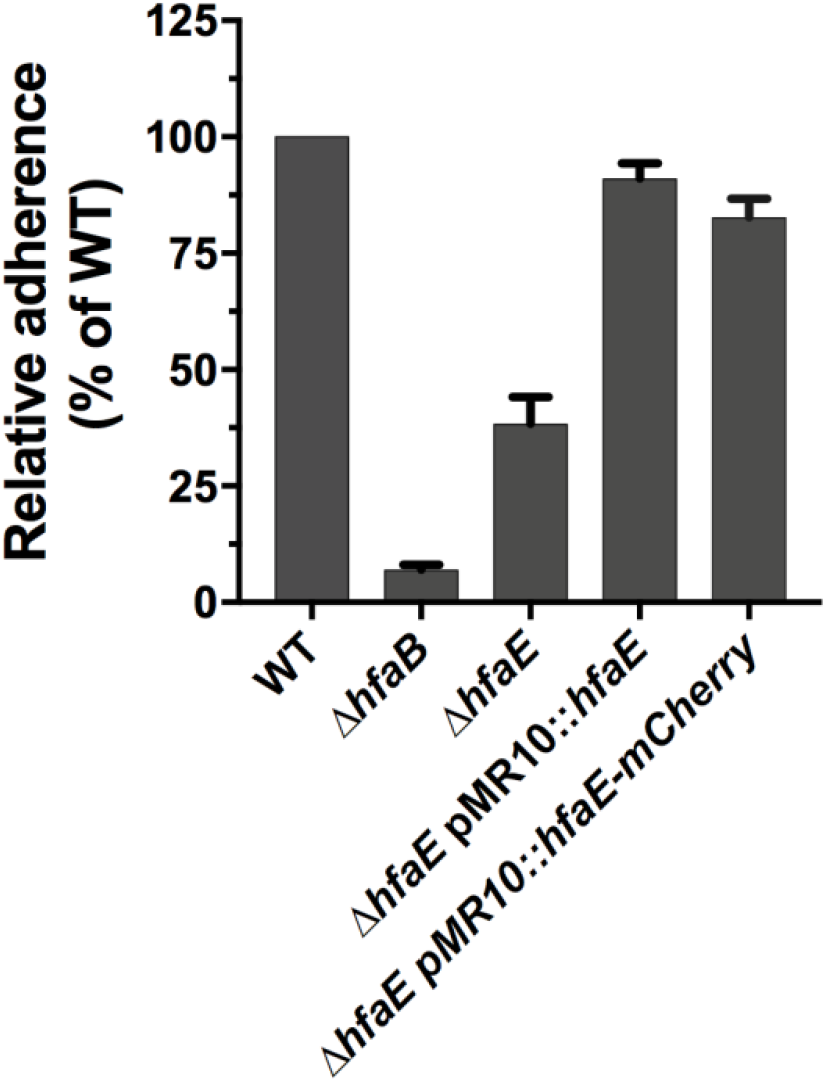
Complementation of *C. crescentus* Δ*hfaE* with *hfaE::mCherr*y, related to figure 4. Quantification of biofilm formation by the crystal violet assay after incubation for 12 h, expressed as a mean percent of WT crystal violet staining normalized to OD_600_. Error is expressed as the standard error of the mean of three independent biological replicates, each with four technical replicates.

**Figure S3:**
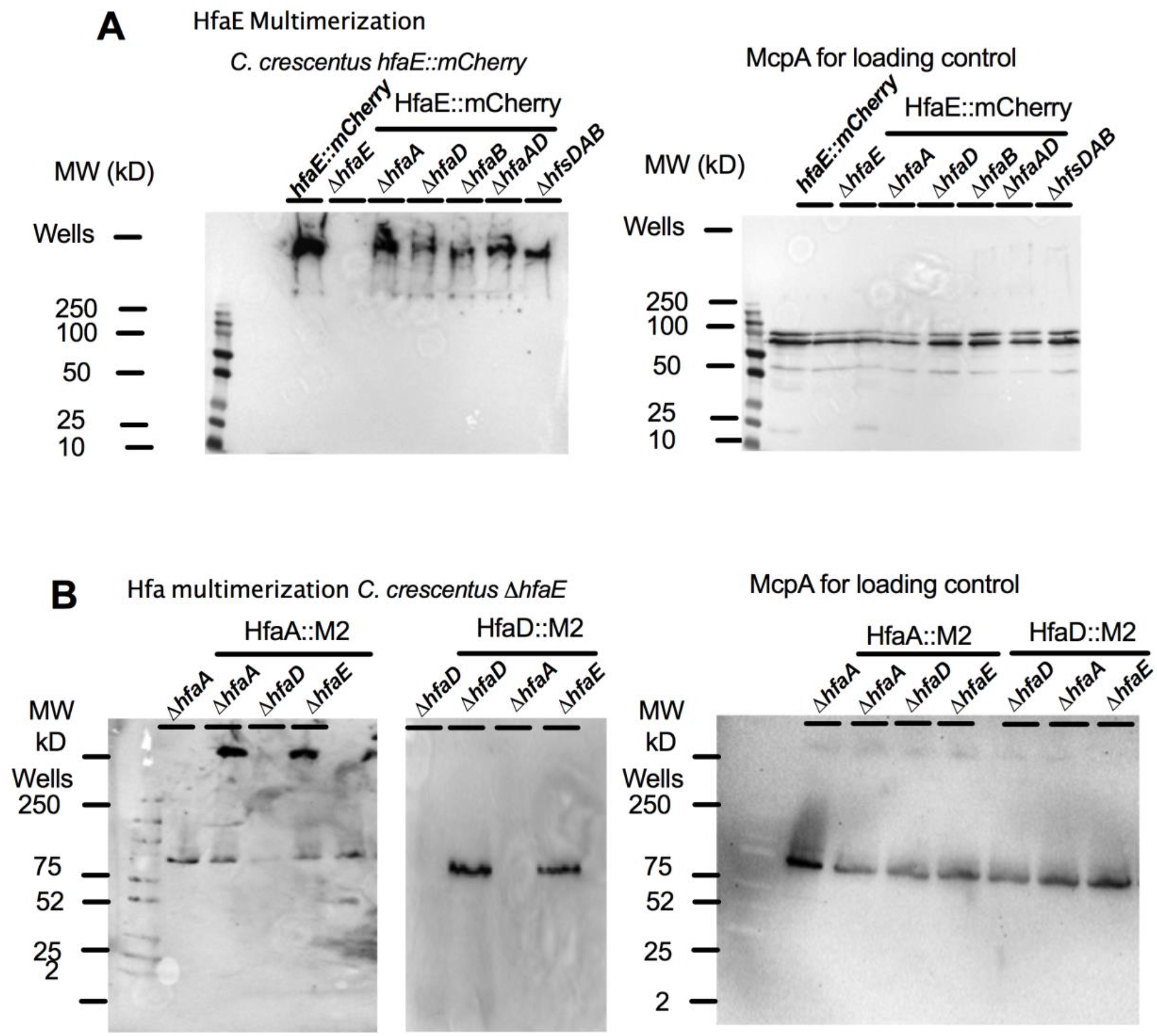
HfaE forms high molecular weight complexes, related to figure 6. **A-B**. Western blots of whole cell lysates of *C. crescentus hfaE::mCherry* (A), *C. crescentus hfaA::FLAG* and *C. crescentus hfaD::FLAG* (B) using anti-FLAG tag (α-M2) antibody with goat anti-rabbit HRP conjugated secondary antibody. Exponentially growing cells grown to an OD_600_ of 0.6-0.8 were normalized to an equivalent of 1 mL of culture at an OD_600_ of 1.0, and 10 ul of the cell lysate was loaded into each lane. Anti-McpA was used as a loading control.

